# The Ventral Tectal Longitudinal Column: A Midbrain Nucleus for Modulation of Auditory Processing in the Cochlear Nucleus, Superior Olivary Complex and Inferior Colliculus

**DOI:** 10.1101/2025.06.24.661350

**Authors:** Brett R. Schofield, William A. Noftz, Yoani N. Herrera, Michael T. Roberts

## Abstract

A ventral tectal longitudinal column (TLCv) has been described in rats and is hypothesized to provide multisensory modulation of acoustic processing in the superior olivary complex (Saldaña et al., 2007, J Neurosci 27, 13108–16). The TLCv is a column of cells in the dorsomedial tectum extending rostro-caudally through the inferior and superior colliculi. It receives ascending auditory input and projects to the superior olivary complex. Further insight into TLCv function has been hampered by limited information on its connections. Here, we provide evidence that a TLCv is recognizable in mice and that it has more extensive connections than previously believed. Deposit of retrograde tracer into the superior olivary complex labels cells bilaterally in the TLCv, comparable to results seen in rats. Viral labeling of neuronal projections demonstrate input to the TLCv from the superior olivary complex and from the inferior colliculus. Thus, the TLCv in mice has inputs and outputs similar to those described in rats. Additional experiments with retrograde tracers revealed more extensive outputs from the TLCv. Neurons in the TLCv are labeled after deposit of retrograde tracers into the cochlear nucleus or into the inferior colliculus. The projections from the TLCv to these targets, like those to the superior olivary complex, are bilateral. These projections are much broader than those described previously. The results suggest that the TLCv could exert modulation over a wide expanse of the auditory brainstem, from the cochlear nucleus through the inferior colliculus.

## 1. INTRODUCTION

The ventral tectal longitudinal column (TLCv) is a midbrain nucleus hypothesized to provide descending modulation of acoustic processing in the superior olivary complex (Saldaña et al., 2007; Aparicio and Saldaña, 2014). The TLCv is a column of cells extending through the inferior colliculus and superior colliculus adjacent to the midline and located just dorsal to the periaqueductal gray (Saldaña et al., 2007). This column of cells stands out after deposit of a retrograde tracer in the superior olivary complex. Faye Lund (1986) placed a retrograde tracer in the superior olivary complex and described that “…a distinct group of labeled cells occurred rostral to the dorsal cortex in the commissure of inferior colliculus, close to the midline…” (p. 42, Faye Lund, 1986). Mulders and Robertson (2001) were the first to describe this group of cells extending rostrally beyond the inferior colliculus; they noted after a deposit of retrograde tracer in the superior olivary complex: “A cluster of labelled neurones was also observed in the commissure of the inferior colliculus, ipsilateral to the injection site… This cluster appeared to spread out in a rostral direction as a continuous band into the superior colliculus” (p. 317, Mulders and Robertson, 2001). Saldaña et al. (2007) labeled this cell column the “tectal longitudinal column” and subsequently designated it the ventral tectal longitudinal column to distinguish it from a dorsal tectal longitudinal column present in the superior colliculus (Aparicio and Saldaña, 2014). In rats, the TLCv comprises ∼11,500 cells, making this nucleus comparable in size or larger than many other brainstem auditory nuclei (e.g., in rats: 9000-10,560 neurons in dorsal cochlear nucleus; 2700-7400 neurons in medial nucleus of trapezoid body; ∼1500-8000 in lateral superior olivary nucleus; Kulesza et al, 2002). In addition to its projection to the superior olivary complex, the TLCv receives input from the superior olivary complex and from the inferior colliculus, and its cells respond to acoustic stimuli (Marshall et al., 2008; Aparicio et al., 2010; Viñuela et al., 2011). Saldaña et al. (2007) provide evidence from Nissl-stained tissue that a TLCv can be distinguished in mouse, guinea pig, hamster, gerbil, chinchilla, rabbit, cat, ferret, macaque monkey and human, suggesting a role in auditory processing across mammals.

Previously, the only known target of TLCv projections was the superior olivary complex, a brainstem center that plays multiple roles in auditory processing. The superior olivary complex is a key center for binaural and temporal processing (Yin et al., 2019). It is also the source of olivocochlear projections, providing the central nervous system with critical controls over cochlear functions (Elgoyhen et al., 2018; Romero and Trussell, 2022). Recent studies highlight roles for modulation of superior olivary complex processing by a number of auditory and non-auditory inputs (e.g., Nevue et al., 2016; Felix et al., 2019; Beebe et al., 2021). TLCv axons terminate in the superior paraolivary nucleus and the ventral nucleus of the trapezoid body and thus could affect many superior olivary complex functions. However, the connectional and physiological data on the TLCv have been obtained only in rats, impeding broader speculation on the functions of the TLCv.

In the course of studying connections among brainstem nuclei, we have acquired numerous observations supporting the existence of a TLCv in mice. In similarity with rats, the mouse TLCv projects to the superior olivary complex and receives input from both the superior olivary complex and the inferior colliculus. We discovered additional projections from the TLCv to the cochlear nucleus and to the inferior colliculus, pathways not previously described in any species. We conclude that the TLCv is present in mice and has connections with the cochlear nucleus, superior olivary complex and inferior colliculus. These properties suggest a much broader role for the TLCv than previously suggested. The TLCv is in a position to modulate auditory processing at multiple brainstem levels, from the cochlear nucleus to the inferior colliculus.

## 2. MATERIALS AND METHODS

All procedures are consistent with NIH guidelines and were approved by the Northeast Ohio Medical University Institutional Animal Care and Use Committee or the University of Michigan Institutional Animal Care and Use Committee. Efforts were made to minimize pain and the number of animals needed for the study.

### 2.1. Animals and tracer deposits

#### 2.1.1. Mouse strains

A total of 39 mice were used (22 females, 17 males). Four mice were C57BL/6J mice (JAX stock no. 000664), in which case experiments were performed at ages preceding high-frequency hearing loss (before P70, Zheng et al., 1999). Four additional mice were VGAT-Cre mice (B6J.129S6(FVB)-*Slc32a1^tm2(cre)Lowl^*/MwarJ); JAX stock no. 028862. The remaining mice were *ChATCre,Cdh23WT* (ChAT-Cre mice), which express Cre recombinase under the choline acetyltransferase, or *ChAT,* promoter (i.e., in cholinergic cells) and carry the wild-type *Cdh23* allele (and therefore do not suffer the early-onset high frequency hearing loss typical of the C57BL/6 J background; Beebe et al., 2020). *ChATCre,Cdh23WT* mice were either heterozygous or homozygous for the *ChAT-Cre* gene but were always homozygous for the wild-type *Cdh23* allele. None of the experiments made use of the Cre expression in the ChAT-Cre or VGAT-Cre animals; rather, these animals were available as part of a breeding program for other experiments and using them here allowed us to minimize use of additional mice. We did not find any differences in the results that could be attributed to the strain, sex or age of the animals.

#### 2.1.2. General surgical procedure

Sterile instruments and aseptic technique were used for all surgical procedures. Animals were anesthetized with isoflurane (2–4% for induction, 1–2% for maintenance) in oxygen. The cornea of each eye was covered with ophthalmic ointment (Moisture Eyes PM, Bausch and Lomb) to prevent drying during anesthesia. The head was shaved and disinfected. Atropine sulfate (0.04 mg/kg, i.p.) was given to some mice to minimize respiratory secretions. The animal was placed in a stereotaxic frame (Leica AngleTwo [RRID: SCR_024708] or Kopf Instruments Model 930-B). Body temperature was maintained with a heating pad. The scalp was incised, the skin retracted and craniotomy was made using a Foredom High Speed Rotary handpiece (Foredom Electric). Retrograde tracer or viral vector was injected with a Nanoliter Injector (World Precision Instruments) or Nanoject III (Drummond Scientific) fitted with a glass micropipette (30-60 μm tip inside diameter) or a 5 μl Hamilton syringe fitted with a 30-gauge needle. The micropipette or microsyringe was oriented vertically or rotated 25 degrees caudally in the parasagittal plane. Following viral vector or tracer deposits, Gel-foam (Harvard Apparatus) was placed in the craniotomy in some mice, and in all mice, the incision was closed with Vetbond adhesive (3M). For some mice, 2% lidocaine hydrochloride jelly (Akorn Inc.) was applied to the incision. All mice received injections of Meloxicam ER (4 mg/kg, s.c., Zoopharm) or carprofen (5 mg/kg, s.c.) to provide extended post-operative pain relief. The animal was placed in a clean cage and monitored until it could walk, eat and drink without difficulty.

#### 2.1.3. Retrograde tracer injections

Retrograde tracers were injected into 3 different groups of mice to label cells that project to the cochlear nucleus, the superior olivary complex or the inferior colliculus. Retrograde tracers were injected into the cochlear nucleus in 8 ChAT-Cre mice (6 females, 2 males; age 4-6 months) and 4 C57BL/6 mice (all females, age < 70 days). Four different retrograde tracers were used: Fast Blue, FluoroGold, green RetroBeads and red RetroBeads (Table 1). Eight of the animals received unilateral deposits into the left or right cochlear nucleus. The remaining 4 mice received bilateral deposits of different tracers into the left and right cochlear nucleus (FB or FG on one side and GB or RB on the other side). Retrograde tracers were injected into the superior olivary complex in four male ChAT-Cre mice (age 9-11 months). Retrograde tracers were injected into the inferior colliculus in 9 ChAT-Cre mice (6 females, 3 males, age 3-8 months).

**Table 1.** Tracers and viral vectors. Summary of concentrations and injection volumes of retrograde tracers and viral vectors used in this study. *from UNC Gene Therapy Vector Core.

| **Retrograde tracer** | **Tracer** | **Concentration** | **Volume injected** |
| --- | --- | --- | --- |
| Cochlear nucleus | Fast Blue | 2.5% in water | 10 nl |
|  | FluoroGold | 4% in water | 50 nl |
|  | Green beads | 100% | 30-100 nl |
|  | Red beads | 100% or 50% in water | 25-100 nl |
| Superior olivary complex | FluoroGold | 4% in water | 15 nl |
|  | Red beads | 100% | 20 nl |
| Inferior colliculus | Fast Blue | 5% in water | 50 nl/site at 1-4 sites |
|  | FluoroGold | 4% in water | 15 nl |
|  | Green beads | 100% or 50% in saline | 50 nl/site at 4 sites |
| **Viral vector** |  |  |  |
| Superior olivary complex | AAV2-hSyn-EYFP* | 5e12 vg/ml | 20-40 nl |
|  | AAV2-hSyn-mCh* | 5e12 vg/ml | 15-30 nl |
| Inferior colliculus | AAV2-hSyn-EYFP | 5e12 vg/ml | 5-40 nl |
|  | AAV2-hSyn-mCh | 5e12 vg/ml | 5-40 nl |

#### 2.1.4. Injection of viral vectors

Two different adeno-associated viral (AAV) vectors were used for labeling neurons in the superior olivary complex or the inferior colliculus (Table 1). These vectors lead to expression of EYFP or mCherry driven by the hSyn promoter, resulting in labeling of neurons in the deposit area and subsequent anterograde labeling of their axons. As noted above, most of the animals used in this study contained Cre-recombinase in a subset of neurons (ChAT-Cre or VGAT-Cre), but the vectors used here were expressed independent of Cre-recombinase. Viral vectors were injected into the superior olivary complex in 3 ChAT-Cre mice (males, 9-10 months old) and 4 VGAT-Cre mice (females, 7-10 months old). In most cases, the vector for EYFP expression was injected into one superior olivary complex and the vector for mCherry expression was injected into the other superior olivary complex, allowing us to evaluate projections from both sides and minimize the number of animals required.

In a separate set of mice, the viral vectors were deposited into the inferior colliculus. As described above for deposits in the superior olivary complex, deposits in the inferior colliculus were also designed with the different vectors injected into the left and right sides of the brain. Vector deposits were made in 7 ChAT-Cre mice (2 females, 5 males, age 5-8 months).

### 2.2. Perfusion and tissue processing

Following time to allow for transport of tracers (5-7 days for retrograde transport; 3-5 weeks for viral experiments), each animal was deeply anesthetized with isoflurane until breathing stopped and corneal and withdrawal reflexes were absent. Then the brain was fixed by transcardial perfusion with Tyrode’s solution or phosphate-buffered saline followed by 50 ml of 4% paraformaldehyde in 0.1M phosphate buffer (PB, pH 7.4) or 10% formalin and 50 ml of the same fixative containing 10% sucrose. Brains were removed and stored overnight at 4°C in the fixative with 25% sucrose. Each brain was frozen and cut into 40 µm thick transverse sections on a sliding microtome. Sections were collected in three series. In some cases, a series was stained for Nissl substance by immersing the sections in NeuroTrace 640/660 (Molecular Probes, Cat. # N2483; dilution 1:100 in PBS) for 20 minutes at room temperature. Individual series of sections could be 1) mounted onto gelatin-coated slides, air-dried, and coverslipped with DPX mountant (Sigma); or 2) placed in freezing buffer (25% glycerol/25% ethylene glycol in 0.1M PB) and stored at -20°C for later processing.

### 2.3. Data analysis and Photography

Fluorescent structures were viewed with a Zeiss AxioImager Z2 microscope equipped with a Hamamatsu Orca camera. For assessing the distribution of labeled cells, the locations of labeled cells in relevant areas were examined at high magnification (40x objective, NA 0.75) and plotted with a Neurolucida system (MBF Bioscience, Williston, VT; RRID SCR_001775) attached to the microscope. Neuronal regions were identified according to a mouse brain atlas (Paxinos and Franklin, 2019) and previous studies (superior olivary complex: Ollo and Schwartz, 1979; inferior colliculus: Choy Buentello et al., 2015; Noftz et al. 2024; TLCv: Saldaña et al. 2007; Viñuela et al. 2011; Aparicio et al., 2014; cochlear nucleus: Beebe et al. 2023). Plots were exported from Neurolucida and prepared for publication with Adobe Illustrator (CC).

Photographs of fluorescent tissue were taken with the Hamamatsu Orca camera and Zeiss AxioImager Z2 microscope controlled either by Neurolucida software (Version 2022, MBF Bioscience) or Zen (Version 3.8, Zeiss) software. Low magnification images were taken with 5x, 10x or 20x objectives and widefield illumination; in select cases the software was used to collect an array of images that were then stitched together. High magnification images were taken with a 63x oil immersion objective (NA 1.4) and structured illumination (Zeiss Apotome 2 or Apotome 3) with optical sectioning (slice depth was set manually at 0.2 μm depth intervals with Neurolucida software or set to optimal depth with Zen software). Final images are maximum projections exported as tif files. Adobe Photoshop CC (RRID SCR_014199) was used to colorize and label images, for global adjustment of brightness and contrast, and for merging fluorescent channels. Image panels were imported into Adobe Illustrator CC (RRID SCR_010279) for final labeling.

## 3 RESULTS

The results comprise 5 series of experiments. We first describe the results from separate retrograde tracing experiments that reveal projections from the TLCv to the cochlear nucleus, inferior colliculus or superior olivary complex. We then describe results from viral deposits in either the superior olivary complex or inferior colliculus. These deposits led to expression of fluorescent proteins that labeled projections to the TLCv via anterograde transport from the deposit sites. In each experimental series, results were similar across sexes, mouse strains and tracers.

### 3.1. The TLCv projects to multiple auditory targets: experiments with retrograde tracers

We deposited retrograde tracers in several different auditory regions to identify cells in the TLCv that project to these regions. We completed three series of experiments associated with deposits in the cochlear nucleus, inferior colliculus or superior olivary complex.

#### 3.1.1. The TLCv projects directly to the cochlear nucleus

Injection of retrograde tracers into the cochlear nucleus labeled cells in many brainstem auditory nuclei. Labeled cells were spread sparsely throughout much of the inferior colliculus, but a collection of cells was notable in the dorsomedial region, adjacent to the midline and just dorsal to the periaqueductal gray. A similar collection of cells was located in the same region throughout the length of the midbrain, extending from the caudal inferior colliculus through the superior colliculus. We obtained similar results with different retrograde tracers, including red RetroBeads (RB), green RetroBeads (GB) and FluoroGold (FG). Figure 1 shows representative tracer deposits from several cases. Figures 1A, B show deposits of FluoroGold that included both dorsal cochlear nucleus and ventral cochlear nucleus. In some cases, the tracer deposit spread into adjacent structures, such as the cerebellum, the trapezoid body, the inferior cerebellar peduncle or the spinal trigeminal tract (e.g., see spread of FG into the cerebellum (Cb) in Fig. 1B). Control injections into these areas outside the cochlear nucleus did not label any cells in the TLCv. Tracer deposits confined to the cochlear nucleus were more readily obtained with RetroBeads. Figures 1C, D show representative deposits of red beads (RB; Fig. 1C) and green beads (GB; Fig. 1D) confined to the ventral cochlear nucleus. Figures 1E-G show cells in the TLCv labeled by the tracer deposits in the cochlear nucleus. The TLCv stood out because of its relatively dense collection of labeled cells that extended rostro-caudally through most of the inferior colliculus and the superior colliculus (Fig. 1H). As noted above, labeled cells were scattered through much of the inferior colliculus, but the collection in TLCv was readily distinguished (e.g., see the most caudal section in Fig. 1H).

**Figure 1.**
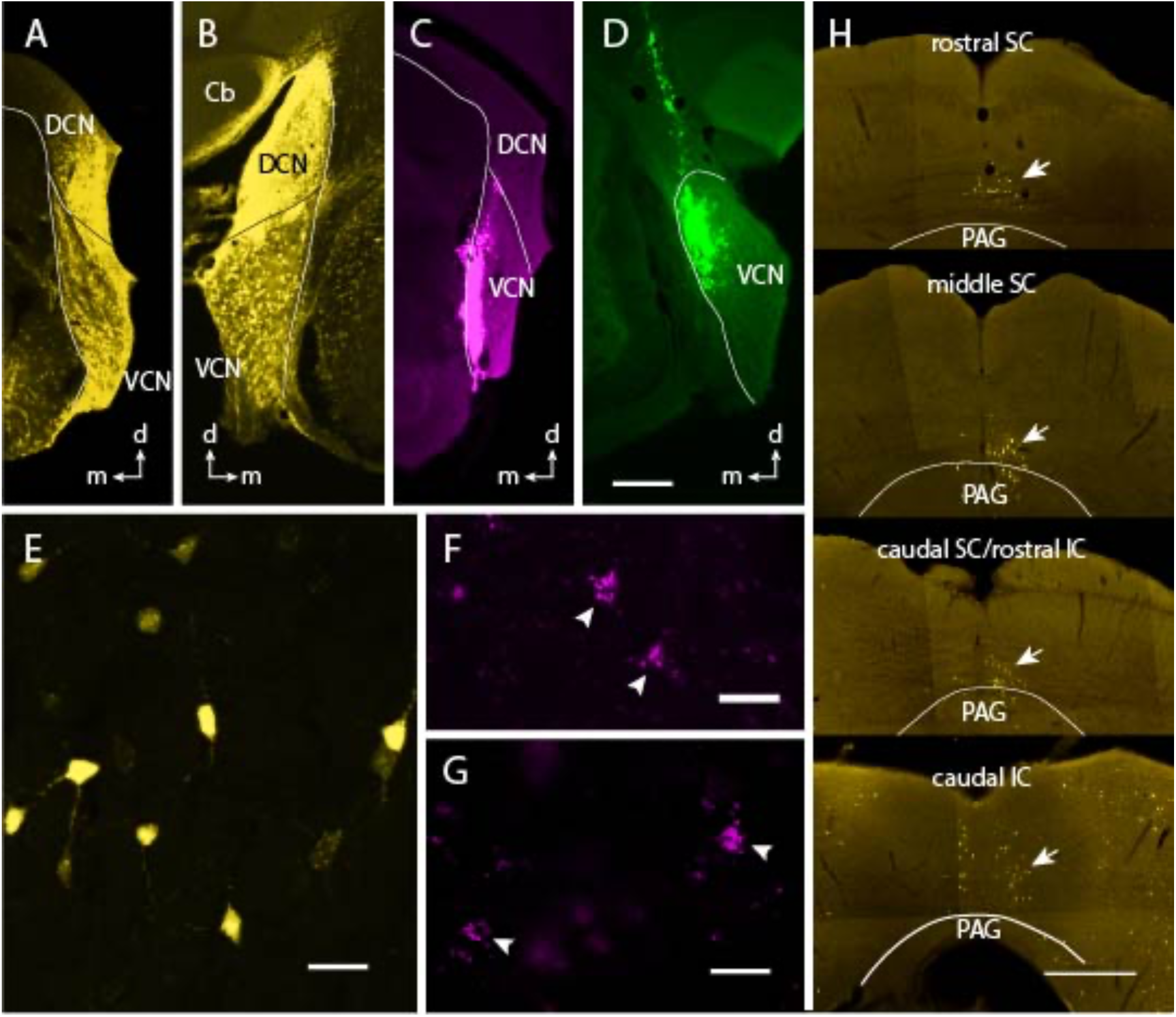
Retrograde tracers show that TLCv projects to the ipsilateral and contralateral cochlear nucleus. **A-D.** Images of representative tracer deposit sites. **A, B**. FluoroGold (FG) deposits that include both the dorsal and ventral cochlear nucleus on the right side in Case M306 (A) and on the left side in Case MVCN1 (B). **C.** Red bead (RB) deposit in the right cochlear nucleus. Case M282. **D.** Green bead (GB) deposit in the right cochlear nucleus. Case MVCN1. **E.** FG-labeled neurons in the ipsilateral TLCv. Case MVCN1. **F, G.** RB-labeled neurons in the ipsilateral TLCv. Case M282. **H.** Labeled neurons in the TLCv were distributed through the rostro-caudal extent of the nucleus. Low magnification images show collections of FG-labeled neurons (arrows) in four sections arranged from rostral (top) to caudal (bottom). Cb, cerebellum; d, dorsal; DCN, dorsal cochlear nucleus; IC, inferior colliculus; m, medial; PAG, periaqueductal gray; SC, superior colliculus; VCN, ventral cochlear nucleus. Scale bars: A-D, H: 0.5 mm; E-G: 20 μm.

We assessed the distribution of TLCv cells that project to the cochlear nucleus by plotting the locations of the labeled cells in every third section through the nucleus. Figure 2 shows the distribution of FG-labeled cells after a deposit of FG in the right cochlear nucleus (deposit shown in Fig. 1A). Labeled cells were plotted in areas adjacent to the TLCv but the figure is limited to TLCv cells to illustrate their rostro-caudal extent and to facilitate comparisons of ipsilateral vs. contralateral sides. FG-labeled cells were found throughout the rostro-caudal length of the TLCv. The ipsilateral TLCv generally contained more labeled cells than the contralateral TLCv, especially at levels of the superior colliculus.

**Figure 2.**
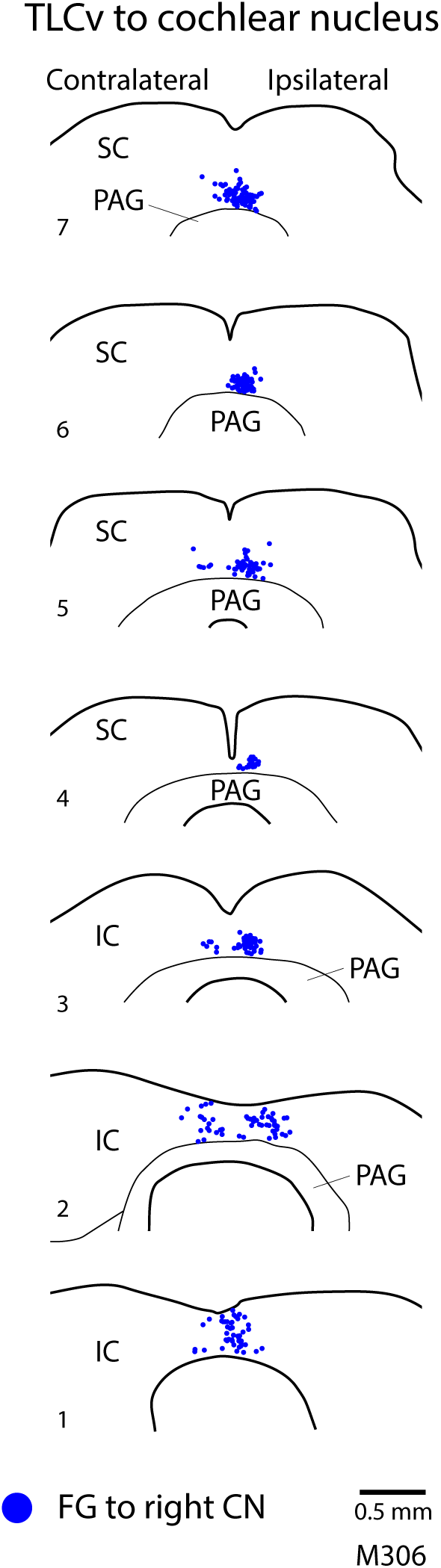
Neurons that project to the cochlear nucleus are found throughout the rostro-caudal length of the TLCv. The figure shows a series of plots of FG-labeled cells (blue circles) in the TLCv after an injection of FG into the right cochlear nucleus (M306; see FG deposit in Fig. 1A). Sections are arranged from caudal (#1 at bottom) to rostral (#7) and are spaced 120 μm apart. Labeled cells were present in the ipsilateral TLCv in all sections. Fewer labeled neurons were present contralaterally.

#### 3.1.2. The TLCv projects directly to the inferior colliculus

In a second series of experiments, we found a projection from the TLCv to the inferior colliculus. Deposits of retrograde tracer (red beads, green beads, FluoroGold or Fast Blue) into one inferior colliculus labeled neurons bilaterally in the TLCv. Figure 3 shows representative results from injections of red beads. Similar results were found with all four tracers. Cells were labeled in the TLCv regardless of the region of the inferior colliculus that was injected. We did not have single deposits confined to the lateral cortex or the intercollicular tegmentum, so we cannot say with certainty that the TLCv targets these two areas, but it appears that that TLCv has fairly broad projections throughout much of the inferior colliculus.

**Figure 3.**
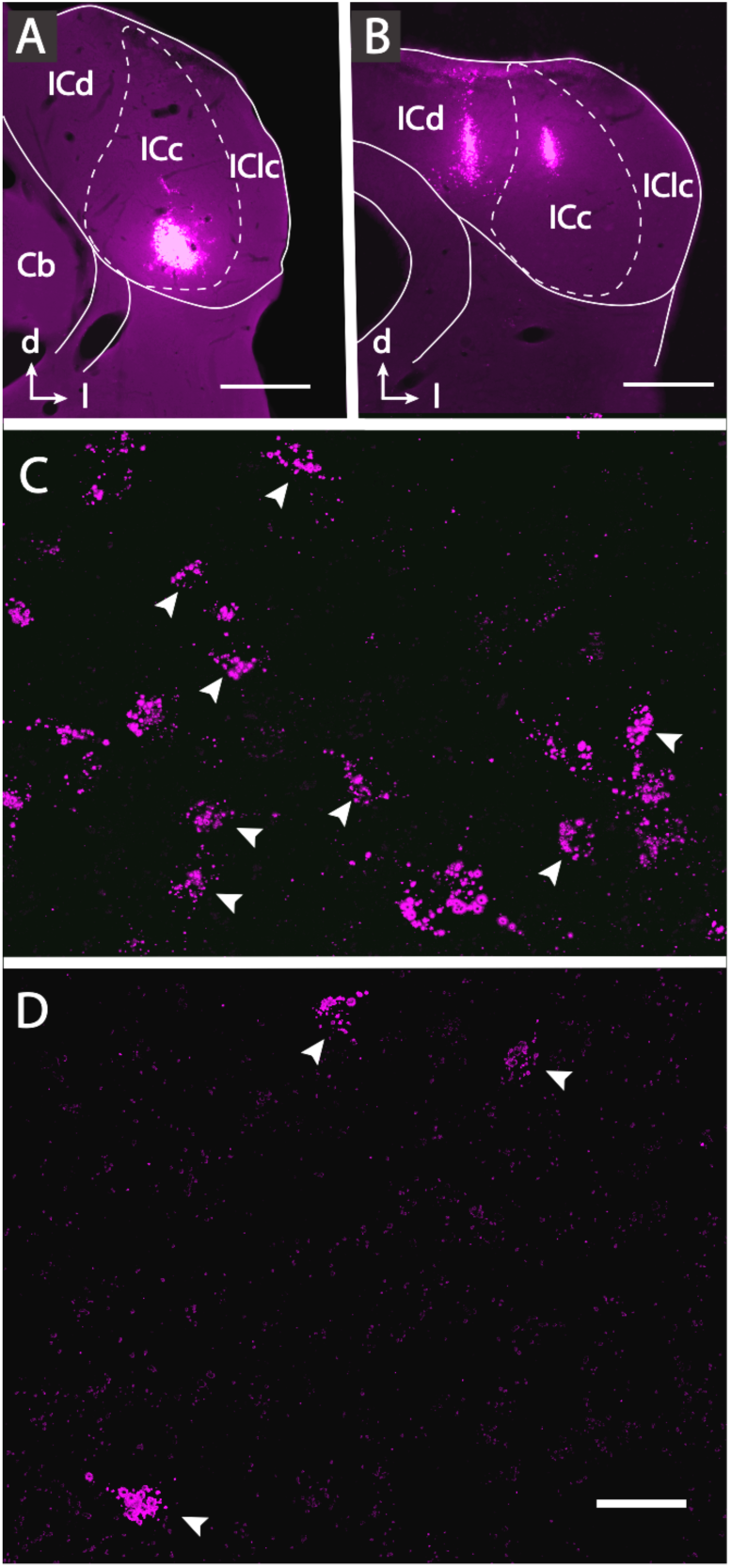
Retrograde tracers show that TLCv projects to the ipsilateral and contralateral inferior colliculus. **A, B.** Images of red bead (RB) deposit sites in two cases. **A.** A single deposit site confined to the ventral part of the central nucleus of the inferior colliculus (ICc). Case M296. **B.** A pair of deposit sites that include the ICc and the dorsal cortex of the inferior colliculus (ICd). Case M218. **C, D.** RB-labeled cells in the ipsilateral TLCv (C) and contralateral (D) TLCv in the case with the tracer deposit shown in panel A. Arrowheads show examples of some of the labeled cells. Case M296. Scale bars: A,B: 0.5 mm; C,D: 20 μm.

In order to assess the distribution of inferior colliculus-projecting cells within the TLCv, we plotted the labeled cells in the TLCv in a representative case (Figure 4). Similar to our results from tracer deposits in the cochlear nucleus, deposits in the inferior colliculus labeled many neurons in the ipsilateral TLCv, throughout its rostro-caudal length. In most cases, sections at the level of the inferior colliculus contained a greater number of cells than those at superior colliculus levels. Labeled neurons were also present contralaterally but were substantially fewer than on the ipsilateral side.

**Figure 4.**
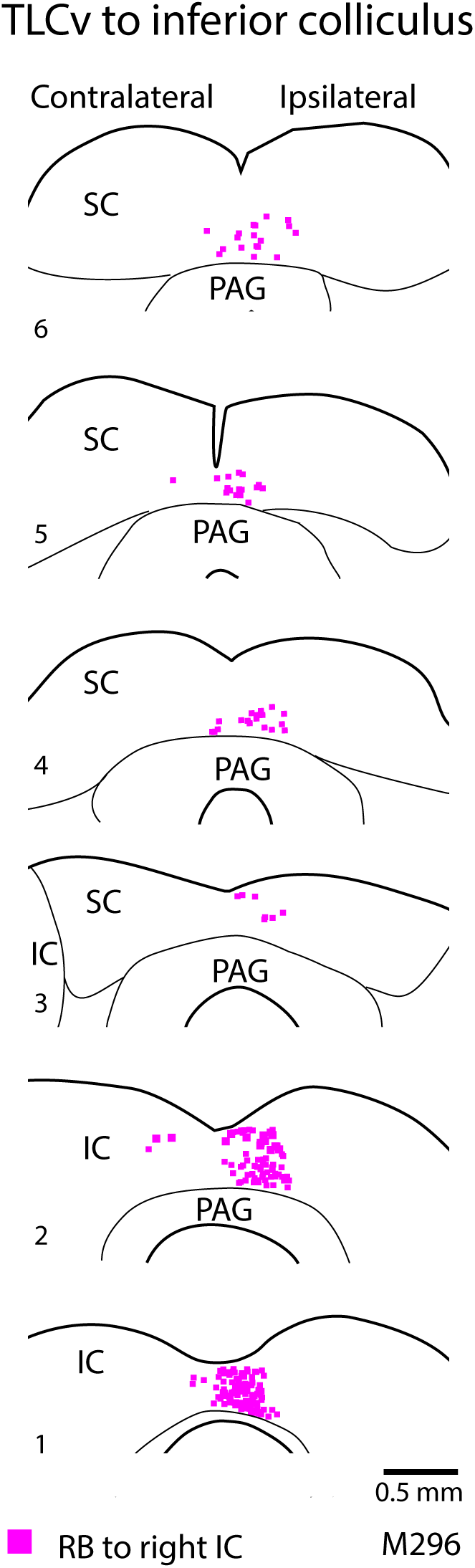
Neurons that project to the inferior colliculus are found throughout the rostro-caudal length of the TLCv. The figure shows a series of plots of labeled cells (magenta squares) in the TLCv after an injection of RB into the right inferior colliculus (M296, tracer deposit shown in Fig. 3A). Sections are arranged from caudal (#1 at bottom) to rostral (#7) and are spaced 120 μm apart. Labeled cells were present in the ipsilateral TLCv in all sections. Fewer labeled neurons were present contralaterally.

#### 3.1.3. The TLCv projects directly to the superior olivary complex

In our third series of experiments, we injected retrograde tracer into the superior olivary complex to look for the pathway from TLCv to the superior olivary complex that has been described in rats (Saldaña et al. 2007). These experiments were based on FluoroGold and RB tracers (7 deposits total; 3 FG and 4 RB). FG diffuses quite readily, allowing a tracer deposit to cover nearly all of the superior olivary complex, whereas the RB tracer typically produces a smaller deposit site. Figure 5 shows examples from a single animal in which RB was deposited in the left superior olivary complex and FG was deposited in the right superior olivary complex. This FG deposit was the largest among all our experiments. It encompassed all the nuclei of the superior olivary complex and spread into the reticular formation dorsal to the superior olivary complex. Each of the FG deposits also spread slightly caudal to the superior olivary complex, encroaching on the lateral paragigantocellular nucleus (a known target of projections from the inferior colliculus; Noftz et al., 2024). None of the deposits, FG or RB, spread rostrally into the adjacent reticular formation or the nuclei of the lateral lemniscus. The FG deposits routinely labeled neurons in many auditory areas (e.g., cochlear nucleus, inferior colliculus); labeled neurons were also present in the TLCv (Fig. 5D). Importantly, similar results were obtained with smaller deposits of RB that were largely confined to the superior olivary complex. Figure 5A shows a deposit of RB centered in the superior paraolivary nucleus that labeled TLCv neurons (Fig. 5C). The primary reason for making bilateral deposits of different tracers in this animal was to minimize the number of animals needed for the experiment. Because the tracers are visualized independently, it also allowed us to examine the tissue for neurons that contained both tracers, indicating that their axon branches to innervate both deposit sites. Figure 5E illustrates 3 RB-labeled neurons in the left TLCv after deposit of RB in the left superior olivary complex. Two of these neurons (arrows) also contained FG, injected into the right superior olivary complex, indicating that they project bilaterally to the left and right superior olivary complex.

**Figure 5.**
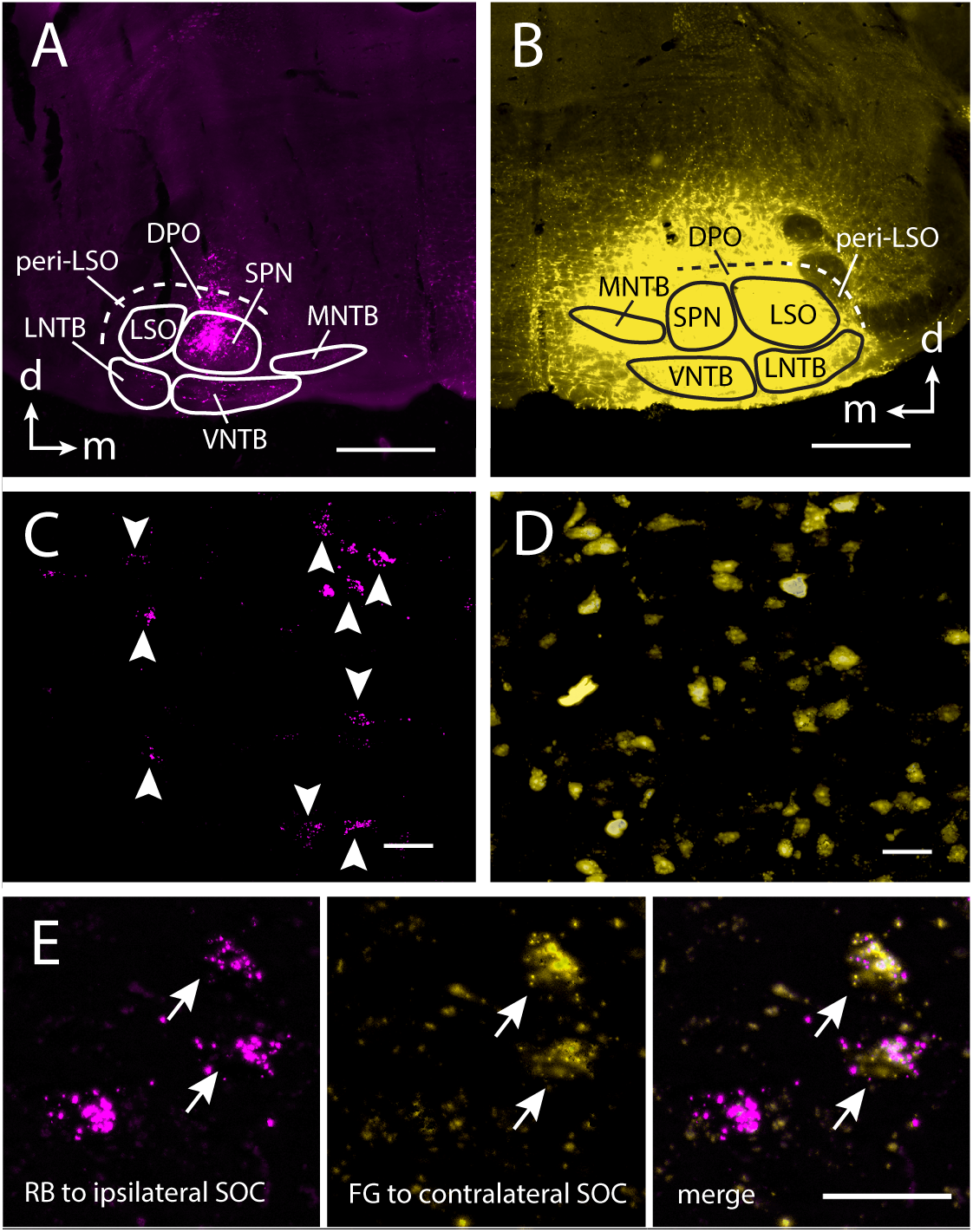
TLCv neurons project to the superior olivary complex ipsilaterally, contralaterally and bilaterally. **A, B.** Tracer deposit sites from a single animal in which RB was deposited in the left superior olivary complex (A) and FG was deposited in the right superior olivary complex (B). Case M354. d – dorsal; m – medial. **C.** RB-labeled neurons (arrowheads) in the left TLCv, ipsilateral to the deposit shown in panel A. **D.** FG labeled neurons in the right TLCv, ipsilateral to the FG deposit shown in panel B. **E.** High magnification image showing labeled neurons in the left TLCv after the deposits illustrated in panels A and B. Left panel: 3 RB labeled neurons; middle panel: same region visualized for FG; right panel: merged image showing that 2 of the neurons were double-labeled with RB and FG (arrows). Scale bars: A, B: 0.5 mm; C-E: 20 μm. DPO – dorsal periolivary nucleus; FG – FluoroGold; LNTB – lateral nucleus of the trapezoid body; LSO – lateral superior olivary nucleus; MNTB – medial nucleus of the trapezoid body; RB – red beads; SPN – superior paraolivary nucleus; VNTB – ventral nucleus of the trapezoid body.

We plotted the locations of TLCv neurons that project to the superior olivary complex to assess their distribution within the nucleus. Figure 6 shows the results from the case with the tracer deposits shown in Figure 5. After a deposit of FG in the right superior olivary complex, retrogradely labeled neurons were present throughout the rostro-caudal extent of the TLCv (Fig. 6A). The neurons were particularly numerous at levels of the inferior colliculus, with substantially fewer labeled cells at superior colliculus levels. Cells were located predominantly ipsilateral to the tracer deposit. This same animal had a deposit of RB in the left superior olivary complex. This deposit was smaller than the FG deposit (compare Figs. 5A vs. 5B) and resulted in fewer labeled neurons in the TLCv. Despite the smaller numbers, the neurons were distributed through most of the rostro-caudal length of the TLCv and were more numerous ipsilaterally than contralaterally (Fig. 6B). In contrast to the FG-labeled neurons, the RB-labeled neurons were more evenly spread between the inferior colliculus and the superior colliculus. As shown in Figure 5E, we were able to identify TLCv cells that contained both RB and FG tracers. These double-labeled cells were also found throughout the length of the TLCv (green stars in Fig. 6C). Because these cells contained both tracers, they are also represented in the FG plot and the RB plot (Figs. 6A and B). To our surprise, the double-labeled cells represented a substantial fraction of the RB-labeled population; Figure 6C highlights this fact with different symbols for the double-labeled cells (green stars) versus cells that contained only RB (magenta squares). In this case, the double labeled cells constituted 25% of the RB cells that projected ipsilaterally and 56% of the RB cells that projected contralaterally. These percentages are based on a small number of cells (57 double labeled cells in a total of 175 cells) from a single case and should not be taken as representative. However, they are notable because the method of double retrograde labeling is biased toward underestimating collateral projections (Schofield et al., 2007). The prevalence of double labeled cells is surprisingly high in this case and suggests that bilateral projections from TLCv neurons to the left and right superior olivary complex are prominent and that future experiments would be warranted.

**Figure 6.**
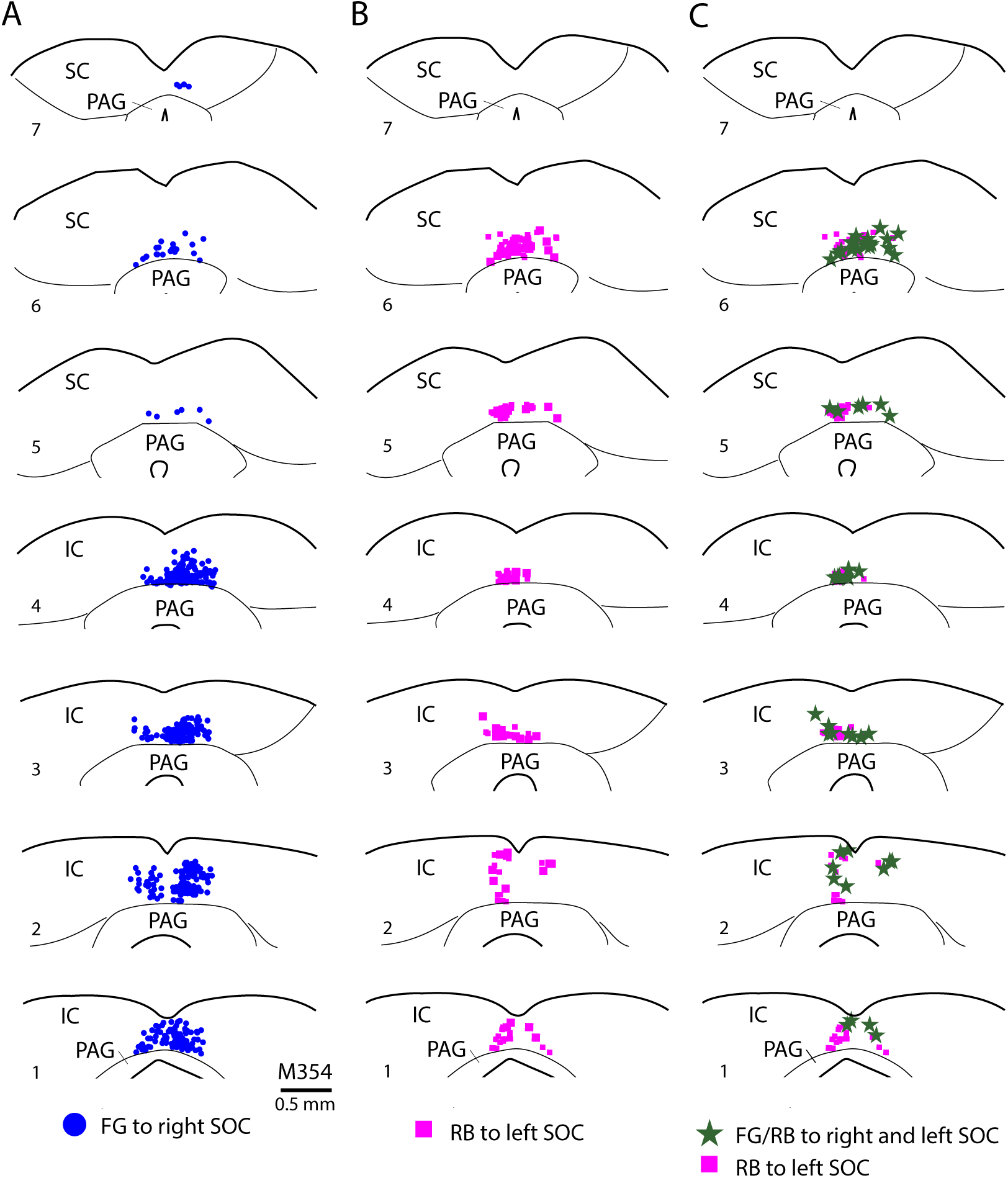
Neurons that project to the superior olivary complex are found throughout the rostro-caudal length of the TLCv. The plots show the distribution of cells labeled after deposits of FG in the right superior olivary complex and RB in the left superior olivary complex (deposit sites shown in Fig. 5A, B). **A, B.** FG-labeled cells that project to the right superior olivary complex are shown in left column A (blue circles) and RB-labeled cells that project to the left superior olivary complex are shown in column B (magenta squares). On both sides of the brain, the TLCv contained both FG-labeled cells and RB-labeled cells. **C.** A substantial number of neurons contained both FG and RB (Column C, green stars). Cells that contained only RB are also shown in column C to illustrate the substantial proportion of RB-labeled cells that were double-labeled with FG.

### 3.2. The TLCv receives input from several auditory regions: Viral labeling of inputs to the TLCv

We used AAV vectors to elicit production of fluorescent proteins in different regions, labeling local neurons as well as their axons. We then assessed the anterograde labeling of the axons to identify projections to the TLCv. We completed two series of experiments, depositing AAV into the superior olivary complex or into the inferior colliculus.

#### 3.2.1. The superior olivary complex projects to the TLCv

We injected viral vectors into the superior olivary complex in 7 animals. In each animal, we directed deposits of AAV2-hSyn-EYFP into the left superior olivary complex and AAV2- hSyn-mCherry into the right superior olivary complex. These vectors lead to constitutive expression (non-dependent on any recombinase) of the fluorescent protein with expression driven by the hSyn promoter to largely limit expression to neurons. Two deposits failed to induce labeling, but ten deposits labeled neurons in the superior olivary complex and, in most cases, some neurons in the reticular formation just dorsal to the superior olivary complex. Two deposits labeled neurons only in the reticular formation dorsal and dorsolateral to the lateral superior olivary nucleus and served as controls; these cases had no anterograde label in the TLCv.

The results were similar across sex, mouse strains and vectors. We describe here the results from a case in which the neuronal labeling was almost entirely confined to the left superior olivary complex. Figure 7A shows the center of the neuronal labeling in case M476, where EYFP-labeled neurons were most numerous in the lateral superior olivary nucleus (LSO), the superior paraolivary nucleus and dorsal periolivary nucleus. Additional neurons were labeled in the peri-LSO region and ventral nucleus of the trapezoid body. No neurons were labeled in the medial nucleus of the trapezoid body or the lateral nucleus of the trapezoid body. EYFP-labeled axons were observed in many known targets of the superior olivary complex, including the cochlear nucleus, nuclei of the lateral lemniscus and inferior colliculus. We confine our description here to the labeled axons in the TLCv. Labeled axons were present throughout the rostro-caudal extent of the TLCv. Figures 7B and C show labeled axons studded with boutons in the ipsilateral TLCv at a mid-rostro-caudal level of the inferior colliculus (Fig. 7B) and at the rostral extreme of the TLCv, at the rostral end of the superior colliculus (Fig. 7C). Axonal branching was prominent, with extensive arbors within the TLCv as well as axons crossing the midline to the arborize on the opposite side. Figures 7D-G show labeled axons at four different rostro-caudal levels. Note first that axons with substantial bouton labeling were present at all levels, and at each level labeled boutons were present bilaterally with a predominance on the ipsilateral side. There was also some variation in the density of axons at different rostro-caudal levels; in the images shown, the densest axons were present at the caudal superior colliculus and caudal inferior colliculus levels (Figs. 7E and 7G). Other cases also showed variations in axon/bouton density, but the exact levels varied, suggesting that different neurons project to different regions (rather than each neuron projecting throughout the TLCv).

**Figure 7.**
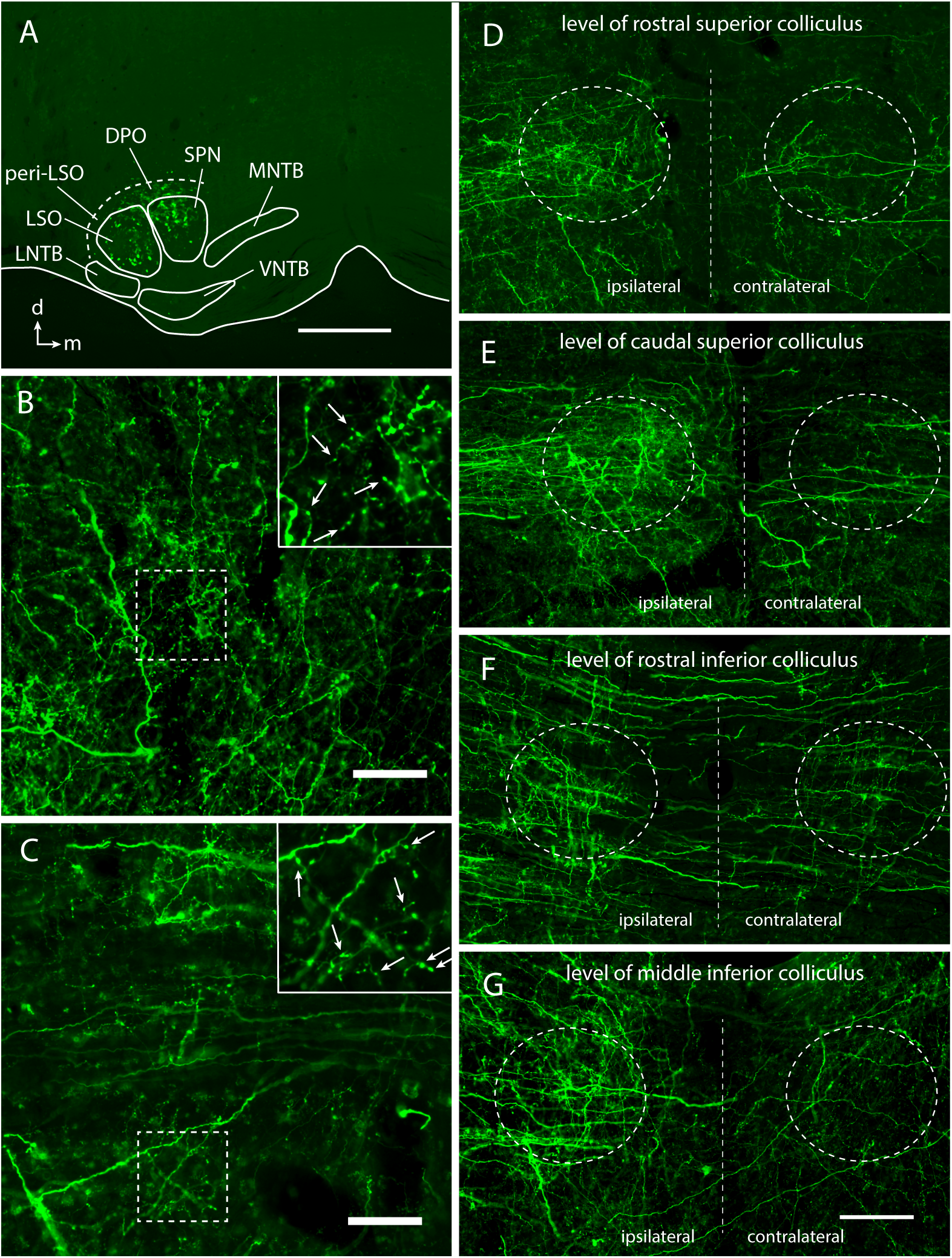
Viral labeling of superior olivary complex neurons labels axons bilaterally in the TLCv. **A.** EYFP-labeled neurons in the left superior olivary complex. Case M476. Scale bar = 0.5 mm. d – dorsal; m – medial. **B-G.** Images showing axons in the TLCv labeled by the viral deposit depicted in panel A. **B, C.** Labeled axons and boutons in the ipsilateral TLCv at the level of the inferior colliculus (B) and rostral superior colliculus (C). Insets show selected regions (dashed boxes) at higher magnification to illustrate labeled boutons (arrows). Scale bars = 50 μm (25 μm for insets). **D-G.** A series of images showing the axonal labeling in the ipsilateral and contralateral TLCv (dashed line ovals) at different rostro-caudal levels of the midbrain (vertical dashed line indicates midline). Scale bar = 100 μm.

As described above, most of our viral deposits labeled neurons in the reticular formation adjacent to the superior olivary complex. Two cases labeled neurons in the reticular formation dorsal to the lateral half of the superior olivary complex but did not label any superior olivary complex neurons. These cases did not label any axons in the TLCv. Several of our deposits labeled neurons in the reticular formation dorsal to the medial half of the superior olivary complex (i.e., dorsal to the superior paraolivary nucleus and the medial nucleus of the trapezoid body) as well as neurons within the superior paraolivary nucleus. Each of these cases had substantial label of axons in the TLCv throughout its rostro-extent and with an ipsilateral dominance. We could not detect any difference due to the inclusion of the reticular formation neurons in these cases versus other cases, so it is possible that the TLCv labeling is entirely attributable to the superior paraolivary nucleus neurons. However, we did not have any cases in which the reticular formation dorsal to the medial half of the superior olivary complex was labeled without involvement of the superior olivary complex. We cannot rule out contributions from these reticular regions, but if they exist, they appear to follow the same patterns as projections from the superior olivary complex.

#### 3.2.2. The inferior colliculus projects to the TLCv

We injected viral vectors to express EYFP or mCherry into the inferior colliculus in seven animals. Four of the animals received dual injections (one for EYFP and one for mCherry expression) at different locations within the same inferior colliculus. None of the deposits labeled neurons in the superior colliculus or the nucleus of the brachium of the inferior colliculus. In two cases, deposits labeled cells in the adjacent cuneiform nucleus and/or periaqueductal gray in addition to cells in the inferior colliculus, so these deposits were excluded, yielding a total of eight viral deposits with labeled cells confined to the inferior colliculus. All remaining cases had labeled axons with boutons distributed bilaterally in the TLCv. Figures 8A-D show results from cases with labeled cells in the central nucleus (ICc) or lateral cortex (IClc) (Fig. 8A) or in the ICc and intercollicular tegmentum (ICT) (Fig. 8B). Case M274 also labeled neurons in the dorsal cortex (ICd) caudal to the ICc (not shown). Both cases showed dense labeling of axons with many boutons in the TLCv (Figs. 8C, D). Similar results were present after a deposit that labeled neurons in the caudal ICc and the caudally adjacent part of ICd. Cases with labeled cells confined to the ICc also labeled axons in the TLCv. Figure 8E shows an example case in which deposits of two different vectors labeled nearby clusters of labeled cells in the left ICc. Both clusters labeled axons within the TLCv (Figs. 8F, G). These ICc cases typically labeled smaller clusters of neurons than cases that included the inferior colliculus shell areas, so it is difficult to interpret differences between cases in the density of labeled axons, but our impression is that the ICc projections are less dense (perhaps much less dense) than those from the inferior colliculus lateral cortex or intercollicular tegmentum.

**Figure 8.**
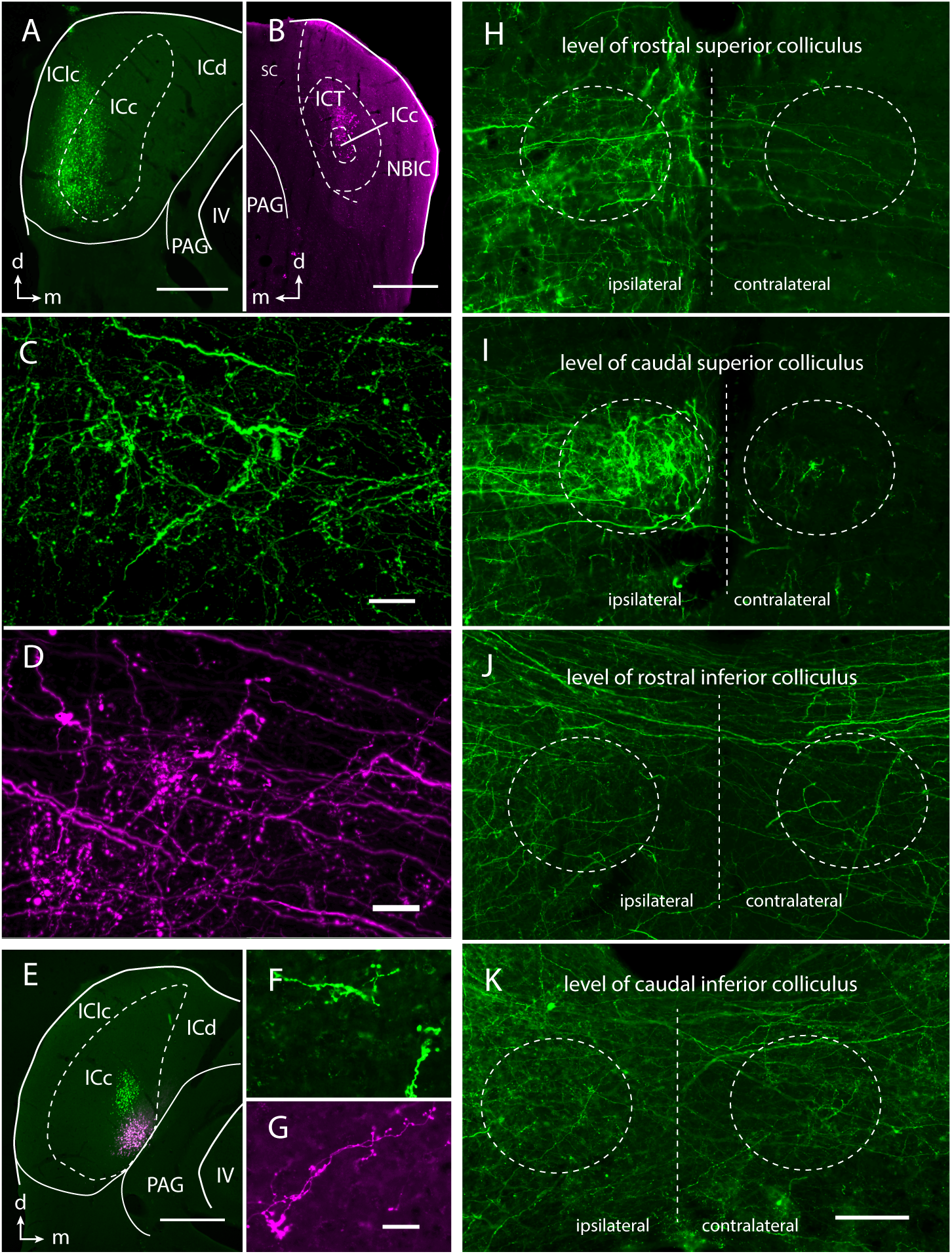
Viral labeling of inferior colliculus neurons labels axons bilaterally in the TLCv. **A, B.** Injections that included inferior colliculus shell regions labeled many axons in the TLCv. **A.** EYFP-labeled neurons in the left ICc and IClc. Scale bar = 0.5 mm. d – dorsal; m – medial. M274. **B.** mCherry-labeled neurons in the rostral part of ICc and adjacent intercollicular tegmentum (ICT). M454. **C, D.** Fluorescent-labeled axons in the ipsilateral TLCv. The panels show results following the viral deposits shown in panels A and B, respectively. Scale bar = 20 μm. **E.** Merged image showing EYFP- or mCherry - labeled cells resulting from separate deposits of virus into the left inferior colliculus. For both deposits, the labeled cells were confined to the ICc. M406. Scale bar = 0.5 mm. **F, G.** Labeled axons and boutons in the ipsilateral TLCv at the level of the inferior colliculus (F) and caudal superior colliculus (G) resulting from the viral deposits shown in panel E. Scale bar = 25 μm. **H-K.** A series of images showing axonal labeling in the ipsilateral and contralateral TLCv (dashed line ovals) at different rostro-caudal levels of the midbrain (vertical dashed line indicates midline) resulting from the viral deposit shown in panel A. Scale bar = 100 μm.

The projections from the inferior colliculus to the TLCv typically exhibited boutons throughout much of the rostro-caudal length of the TLCv. Figures 8H-K show the labeled axons after the viral deposit shown in Figure 8A. As with all other cases, the labeled axons were substantially denser on the ipsilateral side. In this case, the labeling was also much denser in TLCv sections at the level of the superior colliculus than those at the level of the inferior colliculus (compare Fig. 8 panels H, I versus panels J, K). The density of terminations in the rostral (superior colliculus-associated) levels appeared qualitatively similar to that resulting from deposits in the superior olivary complex. In contrast, the inferior colliculus projections to caudal TLCv (i.e., inferior colliculus levels) were noticeably less dense than the projections from the superior olivary complex.

### 3.3 Summary of results

Figure 9 summarizes the results of our tracing studies. Retrograde transport demonstrated direct projections from the TLCv to the cochlear nucleus, the superior olivary complex and the inferior colliculus. The projections to all three targets are bilateral, with ipsilateral dominance.

**Figure 9.**
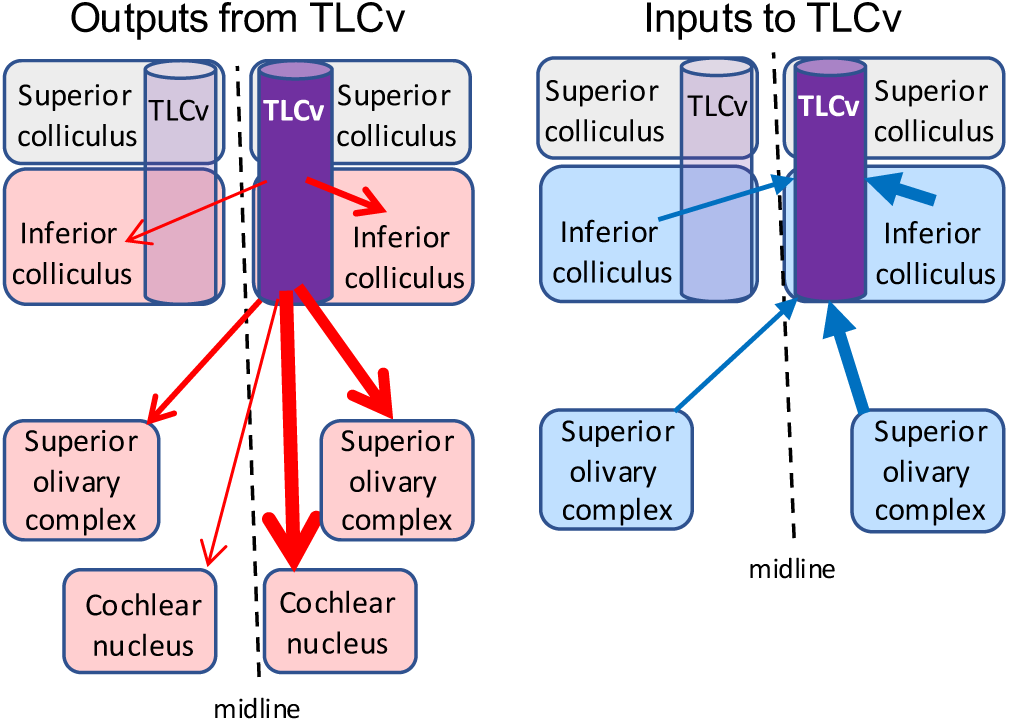
Schematics summarizing the results from the present study. Retrograde tracing revealed bilateral projections from the TLCv to the cochlear nucleus, superior olivary complex and inferior colliculus. Anterograde labeling following viral deposits in the inferior colliculus or the superior olivary complex labeled axons and boutons in the TLCv. Both the inputs and the outputs displayed a bilateral pattern of connections, with ipsilateral components dominant.

Both ipsilateral and contralateral projections originate from most or all of the rostro-caudal length of the TLCv. Viral deposits leading to anterograde labeling demonstrated projections from the superior olivary complex and from the inferior colliculus to the TLCv. These inputs were bilateral with a strong ipsilateral bias. Both sources appear to project through most of the rostro- caudal extent of the TLCv, though there may be differences in the density of terminations at different levels.

## 4. Discussion

Our results demonstrate a column of cells in the dorsal midbrain of mice that have projections to the superior olivary complex and inputs from the superior olivary complex and the inferior colliculus. This column of cells extends rostro-caudally through the levels of the inferior colliculus and the superior colliculus. These properties resemble those described for the TLCv in rats (Saldaña et al. 2007; Aparicio et al. 2010; Viñuela et al. 2011) and support the conclusion that a TLCv is present in mice. In addition, we discovered previously unknown projections from the TLCv to the cochlear nucleus and to the inferior colliculus. The projections to all three targets – cochlear nucleus, superior olivary complex and inferior colliculus – are bilateral with an ipsilateral dominance. These TLCv projections suggest a potentially broad role for the TLCv in modulation of acoustic processing in the auditory brainstem.

### 4.1 Identity of the TLCv

As described in Section 1, Saldaña et al. (2007) suggested on the basis of Nissl stains that a TLCv is present across a wide range of mammals. However, connectional and physiological data on the TLCv have been limited to rats. The present report provides connectional evidence supporting the identification of a TLCv in mice and extends our view of projections from this nucleus.

### 4.2 The rostro-caudal axis of the TLCv

There are numerous properties that display similarities between rostral and caudal parts of the TLCv. The most obvious is the retrograde tracing that first highlighted this cell column (even before its designation as a separate nucleus; Mulders and Robertson, 2001), and that we show here applies to TLCv projections to the cochlear nucleus as well as to the superior olivary complex. The projections from the superior olivary complex to the TLCv also show similarly dense terminations throughout the TLCv rostro-caudal axis. While physiological data are limited to a single study, the results indicate similar responses to acoustic stimuli for cells located in rostral and caudal levels of the TLCv (Marshall et al., 2008).

In contrast, there are also connectional differences that may support different functions for rostral and caudal TLCv. The present results suggest that projections from TLCv to the inferior colliculus may originate primarily from the caudal TLCv (cf. Fig. 4), whereas inferior colliculus inputs to the TLCv appear to terminate more densely in rostral levels (Fig. 8). Another feature of the caudal level of TLCv is a clear overlap with at least part of the nucleus of the commissure of the inferior colliculus. A commissural nucleus, confined to the levels of the inferior colliculus, has been described in several species (e.g., mouse: Meininger et al. 1986; Zettel et al. 1991; Idrizbegovic et al. 1999; rat: Faye Lund and Osen, 1985; Faye Lund 1986; Mulders and Robertson, 2001; rabbit: Herrera et al. 1985; cat: Morest and Oliver, 1984). The commissural nucleus is generally described as extending much more laterally (still within the ICd) than the TLCv, but several of the studies distinguished a medial subdivision of the commissural nucleus that may be closely related to the TLCv. Nonetheless, descriptions of the commissural nucleus have not included any rostral extension into levels of the superior colliculus. Further data are needed to identify possible differences along the rostro-caudal axis of the TLCv and potential relationships with the inferior colliculus commissural nucleus.

### 4.3 Functional implications: descending modulation

Anatomical studies have revealed inputs to the TLCv from the superior olivary complex and from the inferior colliculus (rats: Aparicio et al. 2010; Viñuela et al. 2011; mice: present study) suggesting that TLCv neurons could be driven by sounds. The one available physiological study revealed that TLCv neurons at both inferior colliculus and superior colliculus levels respond to acoustic stimulation, leading the authors to suggest that the TLCv could provide feedback modulation of acoustic processing in the superior olivary complex (Marshall et al. 2008). A primary finding in the present study is the discovery of projections from the TLCv to the inferior colliculus and the cochlear nucleus. Thus, descending modulation could be more widespread than previously suggested, directly affecting neurons bilaterally from cochlear nucleus to the superior olivary complex and the inferior colliculus. An issue that has yet to be explored is the likelihood of multisensory modulation of TLCv activity. Neurons in the TLCv appear to receive inputs from the IClc, an area well known for responses to somatosensory as well as auditory inputs (Aparicio et al. 2010). Whether some TLCv neurons also respond to visual stimuli, as might be expected of neurons in the superior colliculus, is less clear; Marshall et al. (2008) found no TLCv responses to a flash of light, but perhaps other visual stimuli would be effective.

The parabrachial nuclei are considered an integrative zone for interoceptive sensory inputs. These nuclei have widespread projections to many brain areas, including regions from the medulla to the forebrain. Of particular interest to the present study is a group of glutamatergic parabrachial neurons that express neuropeptide S (NPS). Zhang et al. (2024) showed that NPS- expressing parabrachial neurons project to several auditory regions, with particularly distinct projections to the TLCv (see especially their Fig. 11). Interestingly, these parabrachial projections terminate throughout the rostro-caudal length of the TLCv. The functions of the NPS neurons are not known. However, injection of NPS into the brain can cause changes in arousal, body temperature, locomotion and food intake (i.e., functions easily related to integration with interoceptive sensation; reviewed by Zhang et al., 2024). We propose that parabrachial NPS neurons may modulate TLCv neurons in response to interoceptive sensory input.

Interestingly, Zettel et al. (1997) noted that a population of neurons in the nucleus of the commissure of the inferior colliculus in mice immunostain for calbindin or calretinin. The number of these cells can change significantly with age. It appears very likely that these age-related changes included neurons located within the TLCv (see their Figs. 6 and 10). Given the widespread projections of the TLCv, age-related loss of these neurons could affect processing of acoustic stimuli throughout the brainstem.

## 5. Conclusion

We demonstrate that mice have a TLCv with a descending projection to the superior olivary complex similar to the defining characteristic of the TLCv described in rats. Our discovery of TLCv projections to the cochlear nucleus and the inferior colliculus greatly broadens the presumed impact of this nucleus. The presence of ascending input to the TLCv from the superior olivary complex and the inferior colliculus suggests that the TLCv could provide feedback modulation of early auditory processing. The wide divergence of TLCv outputs suggests that feedback from the TLCv could modulate auditory processing at multiple levels of the auditory brainstem, from the cochlear nucleus to the midbrain.

## Author statement

Brett Schofield: Conceptualization, Investigation, Writing - Original draft, Writing - Review & editing, Visualization, Supervision, Funding Acquisition; William Noftz: Investigation, Writing - Review & editing; Yoani Herrera: Investigation, Writing - Review & editing; Michael Roberts: Resources, Writing - Review & editing, Supervision, Funding Acquisition.

## Acknowledgments

This work was supported by the National Institutes of Health grant number NIH R01 DC004391 (to BRS) and NIH R01 DC018284 (to MTR); NIH F31 DC021618 (to Y.N.H.), NIH T32 DC000011 (to Y.N.H.). We thank Nichole Beebe and Michael Kelly for contributions to data generation and Dayanara Lohr and Molly Enrick for comments on an earlier draft of the manuscript.

## Data availability statement

The data that support the findings of this study are available from the corresponding author upon reasonable request.

## Footnotes

1 Abbreviations: AAV, adeno-associated virus; Cb, cerebellum; Cre, Cre-recombinase; ChAT, choline acetyltransferase; DCN, dorsal cochlear nucleus; DPO, dorsal periolivary nucleus; EYFP, enhanced yellow fluorescent protein; FB, fast blue; FG, FluoroGold; GB, Green beads (green RetroBeads); IC, inferior colliculus; ICc, IC central nucleus; ICd, IC dorsal cortex; IClc, IC lateral cortex; ICT, intercollicular tegmentum; LNTB, lateral nucleus of the trapezoid body; LSO, lateral superior olivary nucleus; MNTB, medial nucleus of the trapezoid body; NBIC, nucleus of the brachium of the IC; PAG, periaqueductal gray; RB, red beads (red RetroBeads); SC, superior colliculus; SOC, superior olivary complex; SPN, superior paraolivary nucleus; TLCv, tectal longitudinal column, ventral part; VCN, ventral cochlear nucleus; VGAT, vesicular GABA transporter; VNTB, ventral nucleus of the trapezoid body.

